# Cell-to-cell spread of microsporidia causes *C. elegans* organs to form syncytia

**DOI:** 10.1101/053181

**Authors:** Keir M. Balla, Robert J. Luallen, Malina A. Bakowski, Emily R. Troemel

**Affiliations:** Division of Biological Sciences, Section of Cell and Developmental Biology, University of California, San Diego, 9500 Gilman Drive, La Jolla, CA 92093

## Abstract

The growth of pathogens is dictated by their interactions with the host environment. Many obligate intracellular pathogens undergo several cellular decisions as they progress through their life cycles inside of host cells. We studied this process for several species of microsporidia in the genus *Nematocida* in their co-evolved animal host *Caenorhabditis elegans*. We found that microsporidia can restructure multicellular host tissues into a single contiguous multinucleate cell. In particular, we found that all three *Nematocida* species we studied were able to spread across the cells of *C. elegans* tissues before forming spores, with two species causing syncytial formation in the intestine, and one species causing syncytial formation in the muscle. We also found that the decision to switch from replication to differentiation in *N. parisii* was altered by the density of infection, suggesting that environmental cues influence the dynamics of the pathogen life cycle. These findings show how microsporidia can maximize the use of host space for growth, and that environmental cues in the host can regulate a developmental switch in the pathogen.

## Background

Intracellular pathogens are a diverse category of microbes that rely on the space and resources of their host organisms for replication. After invasion of a single host cell, it is beneficial for intracellular pathogens to spread to other cells to maximize the use of host space for replication before exiting and spreading to new hosts. Several strategies that aid pathogen dissemination within a host have been described for bacterial and viral pathogens. For example, bacterial pathogens in the genera *Listeria, Shigella*, and *Rickettsia* have been shown to utilize host actin to move between host cells by inducing the uptake of a bacterium-containing host cell protrusion into a neighboring cell, thereby avoiding contact with the extracellular space during dissemination [1–3]. Vaccinia viruses have been shown to use host actin in a comparable fashion to spread between host cells [4]. In contrast, several other viruses avoid the extracellular space during spreading by coordinating the fusion of infected host cells with neighboring uninfected host cells to form syncytia [5]. The Gram-negative bacteria *Burkholderia* can also cause host cell fusion as a means for spreading, and fusion is independent of host actin [6]. In addition to spreading through an irreversible fusion of host cells, a recent study found that the bacterial pathogens *Francisella tularensis* and *Salmonella enterica* can be transferred to uninfected cells through an exchange of host cytoplasmic material during partial and temporary fusions of host cells in a process termed trogocytosis [7]. These *in vitro* studies outline a set of growth strategies used by intracellular pathogens to expand their access to host space while remaining inside of host cells, although pathogen dissemination *in vivo* can involve alternative mechanisms. For example, *L. monocytogenes* can cross the barrier of intestinal epithelial cells without host actin via transcytosis into the basal extracellular space [8]. Because animals are composed of diverse and dynamic cells with complex structure, *in vivo* investigations are necessary to understand the relevant mechanisms utilized by intracellular pathogens to grow and spread within the host.

*In vivo* studies are particularly important for studying infection by eukaryotic pathogens, which have especially complex growth dynamics often involving various stages of differentiation that can take place in several specific host environments [9]. Apicomplexan parasites in the genus *Plasmodium*, the causative agents of malaria, require both a Dipteran insect and vertebrate host to complete their life cycle, which has several stages and takes place in many tissues of both hosts. One example of an intracellular strategy used by *Plasmodium* pathogens to spread into uninfected cells involves replicating in large multinucleate structures called merozoites that bud off from liver cells in compartments surrounded by host membrane in transit to a new tissue [10]. Growing and spreading throughout the *Plasmodium* life cycle likely involves several host- and tissue-specific strategies, most of which are not currently understood. Indeed, little is known about dissemination strategies for any eukaryotic pathogen. This gap in our understanding is in part due to the complex nature of the life cycles of eukaryotic pathogens and the host tissues in which they grow.

Microsporidia comprise a large phylum of eukaryotic obligate intracellular pathogens that have multi-stage life cycles. There are more than 1400 species of microsporidia, and they can infect animals ranging from single-celled ciliates to humans [11,12]. The life cycles of microsporidia can be broadly categorized into a replication phase and a spore phase, although various subdivisions of the life cycle have been inferred from diverse morphological data [13]. To initiate infection, a unique microsporidian infection apparatus called a polar tube is thought to pierce the host cell membrane and inject spore contents directly into the host cytoplasm [14]. Replication by many species of microsporidia is carried out in direct contact with the host cytoplasm, and is characterized by nuclear duplication without cell division yielding multinucleate structures called meronts [13]. The final stage of the microsporidia life cycle involves differentiation from meronts into spores, which can exit the host cell and transmit infection. Our understanding of microsporidia growth comes from studies of cells or animals with constant exposure to relatively high levels of pathogen. As such, we currently lack an understanding of the progression that a single microsporidia cell takes as it grows within a single host cell of an intact animal to complete its life cycle.

*Caenorhabditis elegans* is a powerful model system for studying infection by intracellular pathogens in a whole-animal host [15]. Several species of microsporidia in the genus *Nematocida* have been isolated from wild-caught nematodes around the world [16–18]. *Nematocida* species appear to have a life cycle that is generally similar to other microsporidia: after invasion of host cells, *Nematocida* cells replicate in the form of meronts and then differentiate into spores to exit the host cell and propagate infection [17–20]. The best-characterized species is *Nematocida parisii*, which invades and replicates exclusively in the intestine [18]. We recently identified another species of *Nematocida* called *N. displodere*, which invades several tissues including the intestine, muscle, epidermis, and nervous system [17]. *C. elegans* and *Nematocida* species can impart selective pressure on each other, suggesting that these host/pathogen pairs likely coevolve in the wild [21]. The transparency and invariant cellular topology of *C. elegans* together with a collection of their natural microsporidian pathogens provides an ideal system for studying how microsporidia have evolved to grow in dynamic hosts with complex structure.

Here, we characterize the dynamics of the microsporidia life cycle *in vivo* by inoculating *C. elegans* animals with a single pathogen cell in a single host cell. We find that there is a lag phase after invasion, after which growth and replication continues exponentially until sporulation. Surprisingly, *N. parisii* spreads across more than half of the host intestinal cells during the replication phase of growth. By imaging live animals we find that *N. parisii* spreads across host intestinal cell boundaries, causing them to fuse into syncytia that share cytoplasmic space. Growth across host cells before sporulation is a shared property for *Nematocida* species that infect distinct tissues, suggesting that this strategy is conserved for microsporidia. While growth by inducing host cell fusion is shared between *Nematocida* species, there is variation in the tissues that are fused, the speed of growth, and the fitness effects these species have in the host. *N. parisii* and *N*. sp. 1 both grow in the intestine, but *N*. sp. 1 grows faster, spreads more, forms spores sooner, and has a larger negative effect on host fitness than *N. parisii*. Additionally, we find that the pathogen decision to differentiate into spores happens sooner in smaller and denser growth environments. These experiments identify a novel and conserved growth strategy for microsporidia not seen before among eukaryotic pathogens, and illustrate how the host environment influences microsporidia growth and differentiation.

## Results

### Intracellular infection by a single microsporidian cell grows to occupy most of the *C. elegans* intestine

To characterize the *in vivo* growth dynamics of microsporidia, we measured the progression of infection in hosts by generating populations of synchronized infections consisting of a single parasite cell in a single intestinal cell. We pulse-inoculated *C. elegans* with a low dose of *N. parisii* spores to obtain infected populations where most animals were either uninfected, or infected with a single microsporidian cell in a single intestinal cell (Table S1). To visualize infection, we fixed a fraction of the population at various hours post-inoculation (hpi) and stained using a fluorescence *in situ* hybridization (FISH) probe that targets the *N. parisii* small subunit rRNA. At 3 hpi, we found that *N. parisii* cells are small and irregularly shaped, with a single nucleus (Figure 1a). *N. parisii* cells with two nuclei were not observed until 18 hpi (Figure 1b), indicating that replication only occurs after a significant lag time post-infection. By 36 hpi *N. parisii* had grown to spread across several host intestinal cells (Figure 1c). No spores had formed at this time, indicating that *N. parisii* was able to grow across the lateral cell boundaries of neighboring intestinal cells before differentiating into what was previously thought to be the only stage of the microsporidia life cycle able to escape a host cell. New spores were first observed to form at 52 hpi, and by this time *N. parisii* had grown to fill a large fraction of the intestine (Figure 1d). We observed what appeared to be different stages of sporulation, indicated by morphological transitions from tubular meronts to budding rounded structures, rodshaped forms surrounded by diffuse chitin, and smaller rod-shaped cells that stained brightly for chitin, which we loosely define as late meronts, immature spores, and mature spores, respectively (Figure 1e). In addition to defining the transitions that single *N. parisii* cells took through the life cycle as they grew, we observed that meronts grew roughly contiguously in large interconnected structures (Figure 1c-e). We quantified the growth in size of single microsporidia cell infections, and found that initially *N. parisii* took up a miniscule fraction of one *C. elegans* intestinal cell but grew rapidly to take up more than half the space of the entire intestine by the time new spores were formed (Figure 1f).

**Figure 1.**
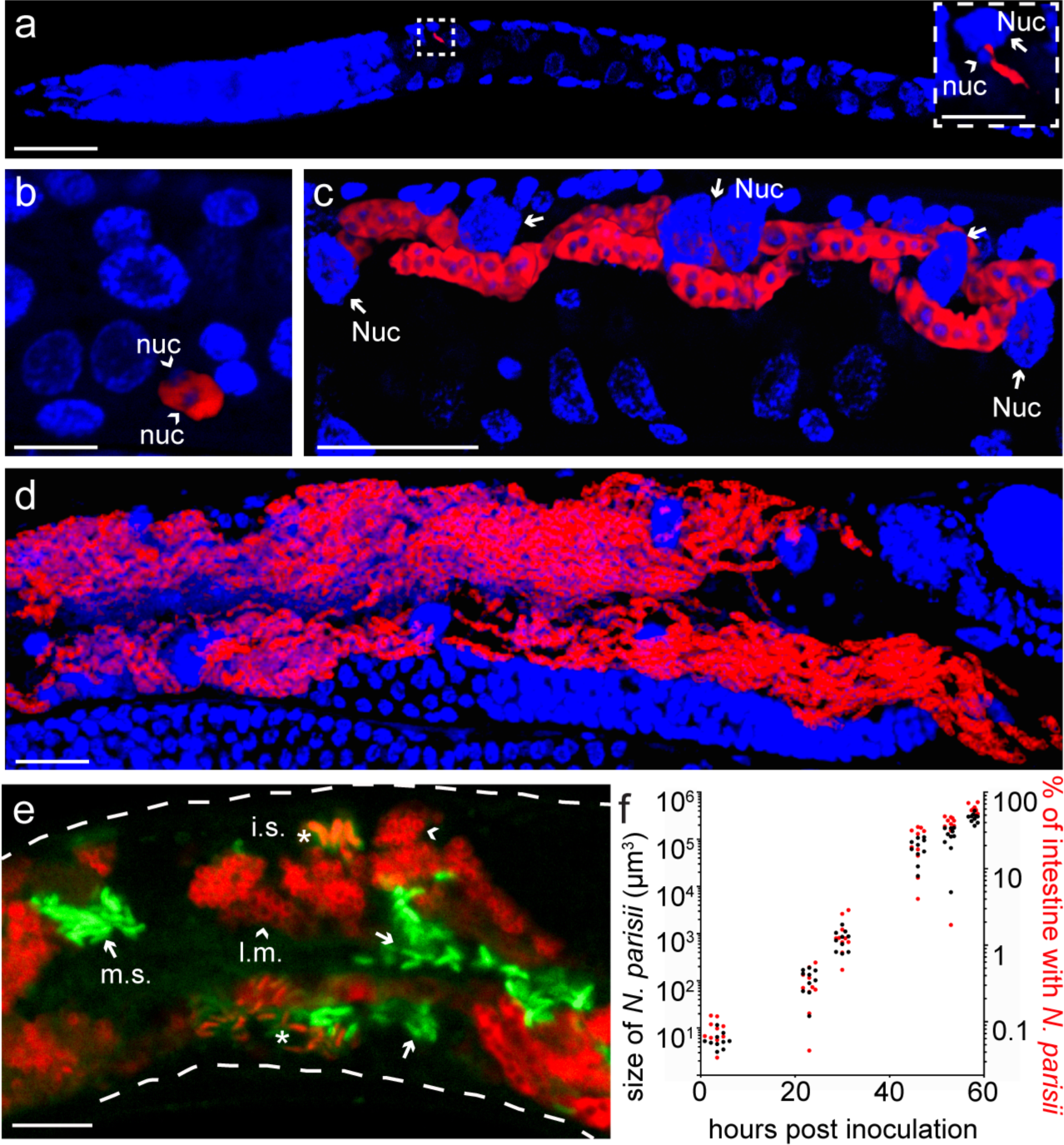
A single *N. parisii* cell can grow to fill most of the *C. elegans* intestine. (A-D) Animals infected by a single *N. parisii* cell, then fixed and stained for DNA with DAPI (blue) and for *N. parisii* with an rRNA FISH probe (red). Images are 3D renderings of confocal z-stacks, and all images are oriented with the anterior of the animal to the left. (A) An animal infected by a single microsporidia cell 3 hpi. Scale bar spans 20 μm. The dashed box encloses a magnified region containing the microsporidia cell and its single nucleus (arrowhead, nuc) next to a larger host nucleus (arrow, Nuc). Scale bar within the box spans 5 μm. (B) Image of infection 18 hpi, marking the beginning of replication by the presence of two pathogen nuclei (arrowheads, nuc). Scale bar spans 5 μm. (C) Image of infection 36 hpi, in which the pathogen has replicated and grown across several intestinal cells (host nuclei indicated by arrows, Nuc). Scale bar spans 20 μm. (D) Image of infection 54 hpi, with extensive growth throughout the intestine and marking the beginning of sporulation. Scale bar spans 20 μm. (E) Magnified image of sporulation 54 hpi in an animal stained with an *N. parisii* rRNA FISH probe (red) and DY96 to label chitin (green). Microsporidia meronts begin to form rounded structures enclosing single nuclei (late meronts indicated by arrowheads, l.m.), rod-shaped cells bearing chitin (immature spores indicated by asterisks, i.s.), and fully formed spores that exclude FISH staining (mature spores indicated by arrows, m.s.). The dashed white line outlines the animal. Scale bar spans 10 μm. (F) Quantification of microsporidia growth from single cells over time. Each dot represents measurement of a single animal. Black dots correspond to measurements of microsporidia volume (left yaxis), and red dots correspond to those same measurements expressed as a fraction of the total intestinal volume (right y-axis).

### *N. parisii* infection spreads across host cell boundaries by fusing neighboring intestinal cells

The growth of single *N. parisii* cells across the host intestine indicated that the boundaries between neighboring host cells might be restructured during infection. To visualize host cell boundaries during single microsporidia cell infections, we infected transgenic animals expressing an intestinal GFP-tagged version of LET-413, a highly conserved protein that localizes to the basolateral membrane of polarized cells [22]. We observed deformation of intestinal cell boundaries as *N. parisii* grew laterally (Figure 2a). Imaging live transgenic animals at this stage of infection demonstrated that *N. parisii* can appose an intact intestinal cell boundary and then grow across that boundary coincident with localized loss of lateral LET-413 (Figure 2b-c, and Video S1). We quantified this growth and found that *N. parisii* infections had spread into one neighboring host cell on average by 27 hpi, and by the time of sporulation the parasite had spread across 12 on average of the 20 total host intestinal cells (Figure S1).

**Figure 2.**
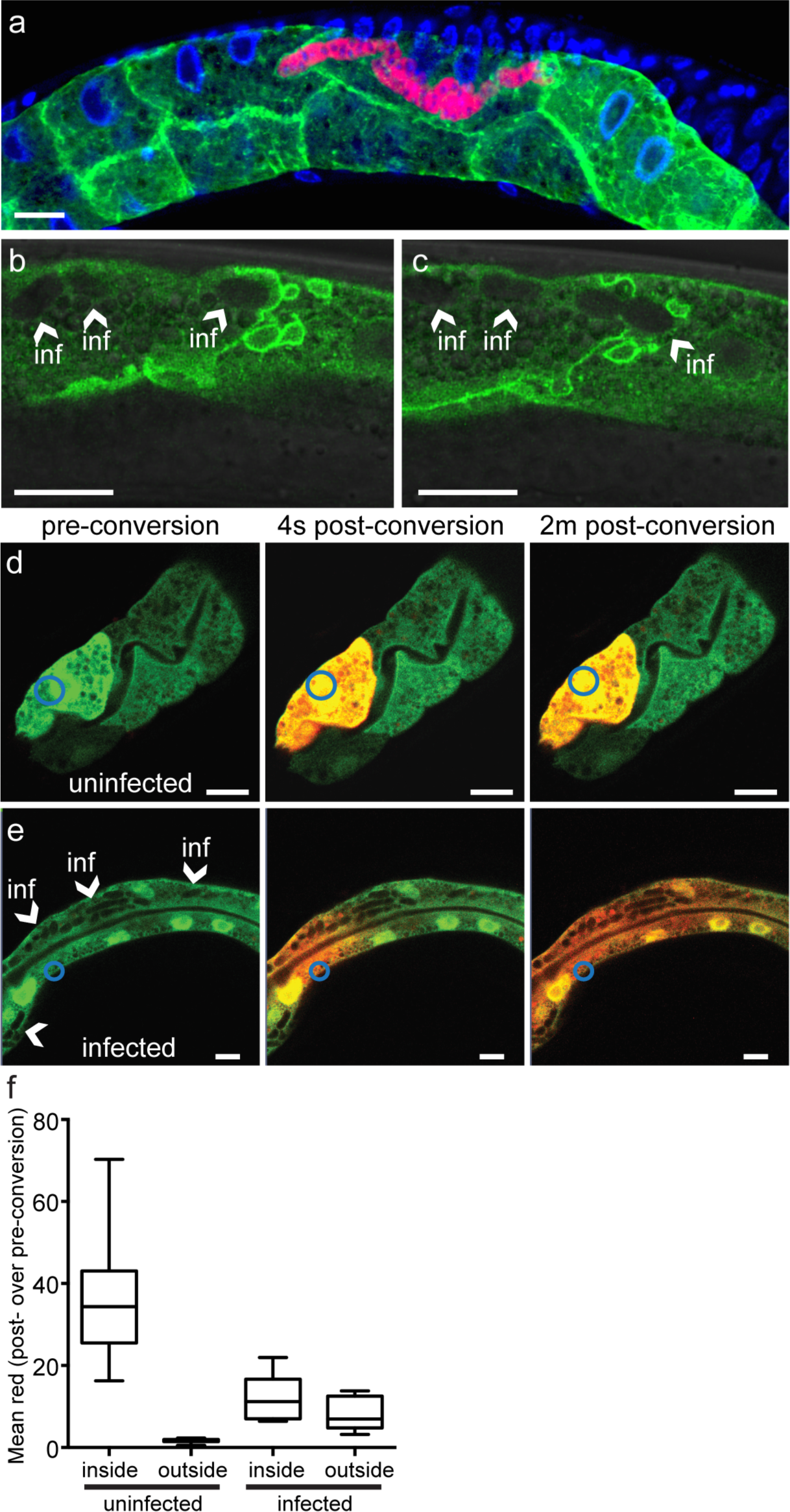
*N. parisii* can spread across and fuse host intestinal cells into a syncytial organ. (A) Image of infection 31 hpi in a fixed transgenic animal expressing a GFP-labeled basolateral protein LET-413 in the intestine (green), stained for DNA with DAPI (blue) and *N. parisii* rRNA with FISH (red). (B) Image of a live infected GFP::LET-413 animal. *N. parisii* is unlabeled but observable as oval-shaped clearings in fluorescent signal (arrowheads). (C) Image of the same animal as in (B) but captured 1 minute later. (D-E) Images of a live uninfected (D) or infected (E) transgenic animal expressing the photoconvertible Dendra protein under an intestinal-specific promoter before conversion (left panel), four seconds after conversion (middle panel), and two minutes after conversion (right panel). Blue circle indicates the region that was targeted for photoconversion. Arrowheads in (E) point to areas in which *N. parisii* can be seen based on the absence of fluorescent signal. Scale bars in all images span 10 μm. (F) Quantification of red Dendra signal diffusion in 10 uninfected cells of infected animals, or 10 infected cells. The amount of red signal was measured before and 2 minutes after conversion within the targeted cell and outside of the targeted cell (>30 μm from cell boundary into neighboring cell). Box plots show the fold change in red signal post- over pre-conversion. The labels for inside and outside indicate the cell that was measured with respect to the cell targeted for photoconversion.

The results above indicate that *N. parisii* spreads across cell boundaries during growth, leading to the question of whether these microsporidia-induced modifications cause neighboring intestinal cells to join together and share cytoplasmic space. To address this question, we generated transgenic animals that express the green-to-red photoconvertible fluorescent Dendra protein in the cytoplasm of their intestinal cells. After infecting these animals, we photoconverted Dendra in a single cell from green to red and then recorded the diffusion of the red signal. When we photoconverted uninfected cells within an infected animal, we found that the red signal remained restricted to the cell in which it was converted (Figure 2d), indicating that the boundaries of host intestinal cells not yet reached by *N. parisii* were intact. In contrast, when Dendra was photoconverted in an infected cell, we found that the red signal diffused across all neighboring cells where infection was present (Figure 2e). When quantifying this diffusion we found little red signal outside of photoconverted uninfected cells, but large amounts of red signal outside of photoconverted infected cells (Figure 2f). These data show that *N. parisii* is able to spread across the boundaries of neighboring host cells, and in so doing cause intestinal cells to join together into syncytia and share cytoplasmic space. Thus, we show that a eukaryotic pathogen causes what appear to be abnormal cell fusion events in the host to allow for intercellular spread.

### Infection by different *Nematocida* species causes syncytia formation within distinct host tissues

Microsporidia species vary in the host tissues where they can infect and replicate. We investigated whether infection-induced fusion of host cells is a conserved growth strategy for microsporidia by characterizing the growth of two other *Nematocida* species from single cells. Like *N. parisii, N*. sp. 1 only invades and replicates in the intestine of *C. elegans*, while *N. displodere* invades and replicates within several tissues [17,18]. To compare the ability of different microsporidia species to cause host-tissue syncytia formation, we performed single-cell infections of animals with these three *Nematocida* species (Table S1) and quantified the number of host cells with infection after growth to the point of spore formation. Of note, these experiments were performed at 15°C to facilitate *N. displodere* replication, while other experiments were performed at 20°C. First we analyzed infections in the intestine, and here we found that both *N. parisii* and *N*. sp. 1 always spread across several host intestinal cells by the time of spore formation (76 hpi), but that *N. displodere* was most often restricted to a single intestinal cell at its time of spore formation (120 hpi) (Figure 3a-c,e). Next, we analyzed spread in the muscle. Unlike skeletal and somatic muscle cells in other animals, the 95 body wall muscle cells of *C. elegans* do not fuse into syncytia during normal development [23]. However, we observed that single-cell *N. displodere* infections grew across many host muscle cells before forming spores, while *N. parisii* and *N*. sp. 1 did not invade or replicate in the muscle (0/60 animals analyzed) (Figure 3D,E). Furthermore, we observed four cases of single *N. displodere* cell infections that appeared to have spread out of the large hypodermal syncytium of *C. elegans* (hyp7) and into the anterior epidermal cells (Figure S2) [24]. These data demonstrate that at least among natural pathogens of nematodes, intercellular spread through host cell syncytia formation is a conserved growth strategy for microsporidia determined by species- and tissue-specific interactions.

**Figure 3.**
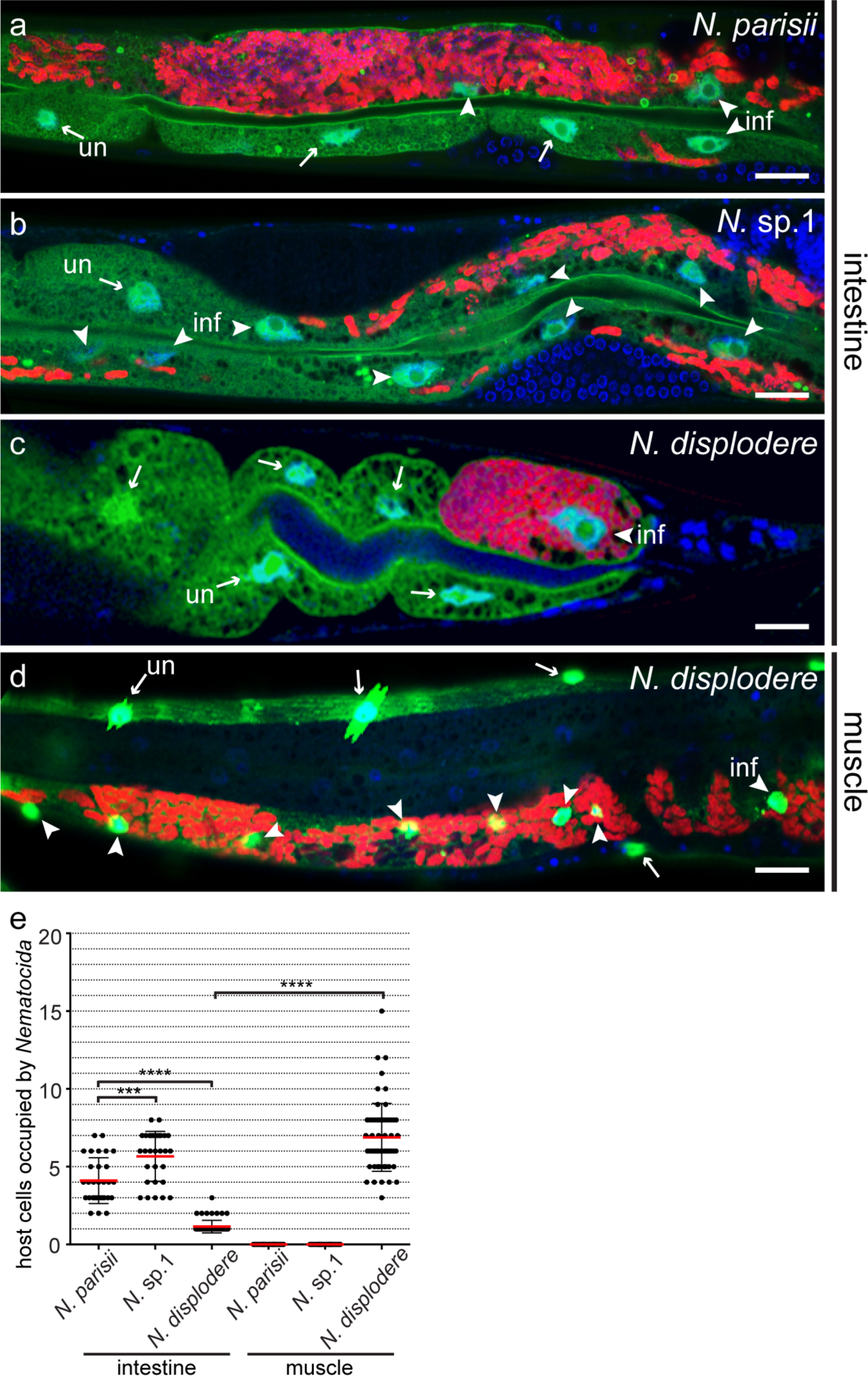
Spreading across host cells is a conserved microsporidia growth strategy with distinct host cell fusion patterns caused by distantly related *Nematocida* species. Images of single-cell infections by *N. parisii* (A), *N*. sp. 1 (B), or *N. displodere* (C) in the intestine at the time of sporulation. Transgenic animals with GFP expressed in the cytoplasm and nuclei of intestinal cells (green) were fixed and stained for DNA with DAPI (blue) and *Nematocida* rRNA with FISH (red). Arrowheads point to host nuclei of cells with infection (inf), arrows point to host nuclei of cells without infection (un). Scale bars in all images span 20 μm. (D) Image of a single-cell infection by *N. displodere* in a transgenic animal with GFP expressed in the cytoplasm and nuclei of muscle cells. (E) Quantification of the number of host cells occupied by parasite by the time of sporulation (76 hpi for *N. parisii* and *N. displodere*, 120 hpi for *N. displodere*, all at 15°C). Data are combined from two independent experiments with at least 15 animals measured per condition. Each dot is a measurement from a single animal. Red bars show averages, black bars show standard deviations. Significance was tested by one-way ANOVA, p-values indicated by asterisks with ***≤0.001, and ****≤0.0001.

### Microsporidia species vary in their intestinal growth dynamics and effects on host fitness

While both *N. parisii* and *N*. sp. 1 grow in the intestine of *C. elegans*, we found that single *N*. sp. 1 cells spread into more host intestinal cells on average than single *N. parisii* cells by the time of spore formation (Figure 3e). This observation indicated that there could be differences in the dynamics of growth between species of microsporidia despite the fact that they start off in the same environment. To compare the dynamics of intestinal pathogen growth between species of microsporidia, we infected animals with single *N. parisii* or *N*. sp. 1 cells and measured their rates of growth and spore formation over time. Both species grew from a single cell to replicate approximately 50,000 times before forming spores (Figure 4a). Interestingly, *N*. sp. 1 grew slightly faster and formed spores much earlier than *N. parisii*. Rates of growth were non-uniform over the course of infection, but the average doubling times during the exponential stages of growth (18 hpi – 48 hpi) for *N. parisii* and *N*. sp. 1 were 2.4 h and 2.1 h, respectively. In addition to completing its lifecycle more rapidly, *N*. sp. 1 infection had a much stronger negative effect on host fitness than *N. parisii* infection. Single *N. parisii* cell infections slightly reduced the growth of developing animals compared to uninfected animals, while single *N*. sp. 1 cell infections stunted practically all host growth (Figure 4b). Furthermore, *N. parisii* infection decreased host egg production at 60 hpi compared to uninfected animals while *N*. sp. 1 infection essentially eliminated host egg production (Figure 4c). These observations reveal that related microsporidia species can have distinct growth dynamics in the same niche and can differentially exploit host space with varied effects on host fitness.

**Figure 4.**
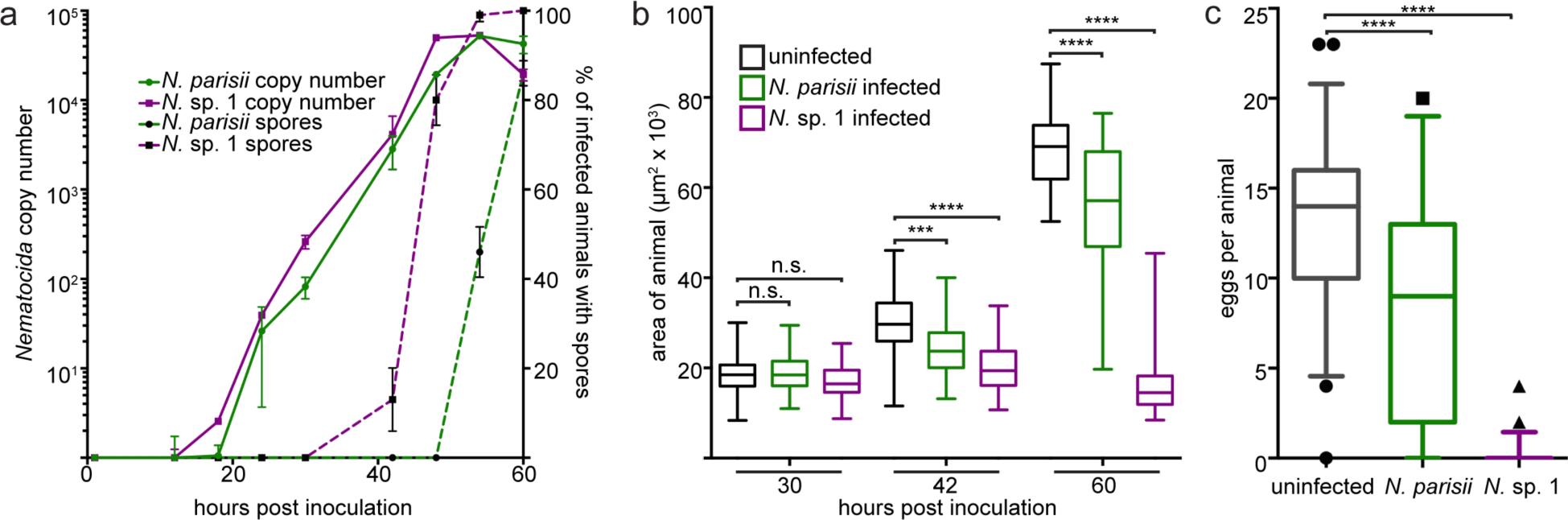
*Nematocida* sp. 1 completes its lifecycle faster and impairs fitness of *C. elegans* more than *N. parisii*. (A) Growth kinetics of single-cell *N. parisii* and *N*. sp. 1 infections. Purple lines correspond to *N*. sp. 1 kinetics, green lines correspond to *N. parisii* kinetics. Solid lines show the kinetics of microsporidia replication, dashed lines show the kinetics of spore formation in the same populations of animals. Averages of two biological replicates are shown with standard deviation. (B) Sizes of animals infected by single *N. parisii* cells (green boxes) or *N*. sp. 1 cells (purple boxes) compared to uninfected animals (black boxes) over time. Box-and-whisker plots include data from 50 animals per condition. (C) Eggs in animals infected by single *N. parisii* cells (green boxes) or *N*. sp. 1 cells (purple boxes) compared to uninfected animals (black boxes) at 60 hpi. Boxand-whisker plots include 95% of the data from measurements of 50 animals per condition; dots represent the remaining 5%. Significance was tested by one-way ANOVA, p-values indicated by asterisks with ***≤0.001, and ****≤0.0001.

### Host body size and infection density affect the timing of microsporidia sporulation

Based on the kinetics of microsporidia growth (Figure 1f and Figure 4a), at least two major transitions are apparent: a lag phase during which pathogen cells are not replicating after invasion of host intestinal cells, and a sporulation phase in which some pathogen cells stop replicating and form new spores to complete the life cycle. We noticed that infection by *N*. sp. 1 caused animals to be smaller than infection by *N. parisii* (Figure 4), and therefore hypothesized that differences in host size might influence the timing of the decision to form new spores. To test this possibility, we infected body size mutants with single *N. parisii* cells and compared spore formation timing to wild-type animals. The *sma-6* and *lon-1* mutations occur in members of the transforming growth factor (TGF) beta signaling pathway that lead to shorter or longer animals, respectively [25,26]. Animal size is positively regulated by *sma-6*, which encodes a TGF beta-like type I receptor, while *lon-1* encodes a downstream factor that is negatively regulated by signaling through SMA-6. Somewhat surprisingly, we found that both the *sma-6* and the *lon-1* mutant strains were smaller in total area than wild-type animals, providing a useful set of strains for testing the hypothesis that body size could affect the growth dynamics of microsporidia (Figure 5a). Interestingly, we found that single pathogen cell infections in these small mutant animals had formed spores earlier than wild-type animals (Figure 5b). These differences could result from a decrease in the time it takes for *N. parisii* to fill the host intestine, potentially providing a density-dependent cue to transition into the spore formation stage of the life cycle. We tested for the influence of infection density on the timing of spore formation by increasing the initial multiplicity of infection, which would increase the rate at which *N. parisii* fills the host tissue. In this experiment, we pulse-infected animals with a range of *N. parisii* spore dosages and measured new spore formation at 52 hpi. Consistent with our hypothesis, we found that more spores formed when we increased the multiplicity of infection (Figure 5c). Because all infected animals will eventually be full of spores, we interpret the increase in spores formed at 52 hpi as evidence that spores form sooner in smaller animals or animals that were infected by multiple microsporidia cells. Furthermore, we found that the growth rate of *N. parisii* was similar in hosts of different sizes and at different multiplicities of infection, demonstrating that infection density affects the timing of spore differentiation, but does not affect the lag phase or rate of replication (Figure S3). Thus, host size and infection density appear to influence the timing of a switch from replication to differentiation into spores for microsporidia.

**Figure 5.**
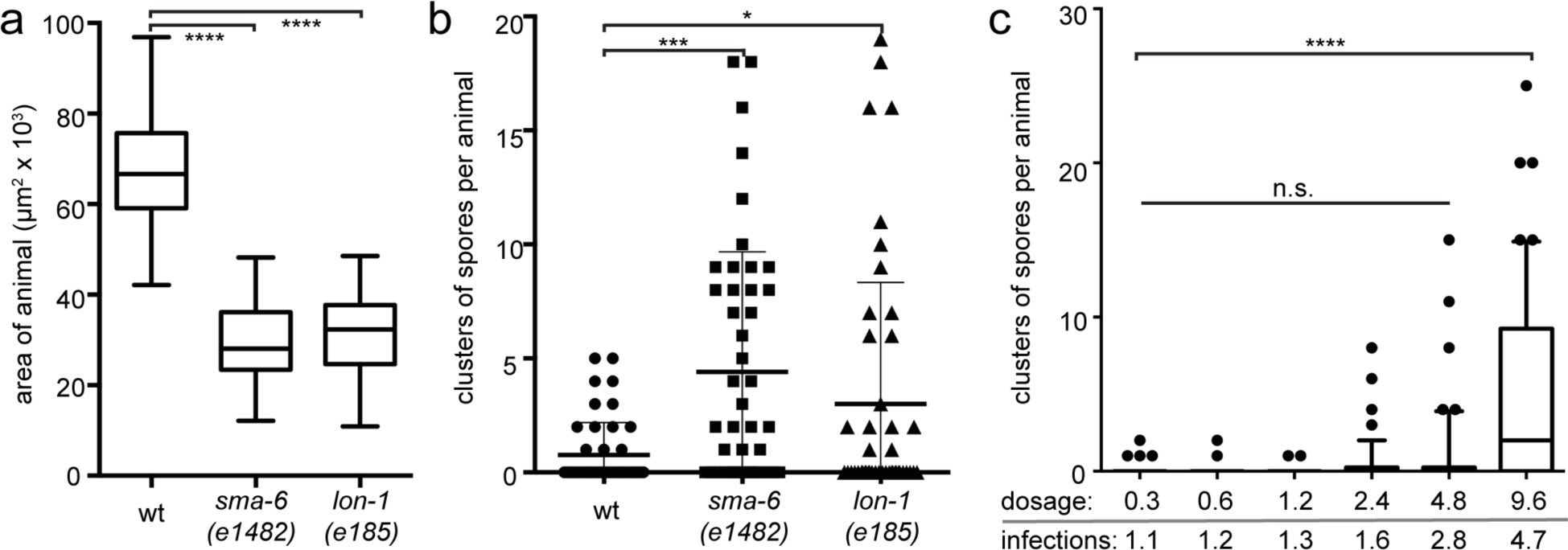
*N. parisii* forms spores faster in denser growth environments. (A) Sizes of wild-type and size mutant animals infected by single *N. parisii* cells 53 hpi. Box-and-whisker plots include data from 50 animals per condition. (B) Spore clusters per animal measured in the same animals as in (A). Dots represent measurements of individual animals; bars show averages with standard deviations. (C) Spore clusters per animal 52 hpi in populations that were pulse-infected with different dosages of *N. parisii* spores. Dosages of spores (x106) are shown underneath the graph, along with the average number of infections per animal at each dosage. Box-and-whisker plots include 95% of the data from measurements of 50 animals per dosage; dots represent the remaining 5%. Significance was tested by one-way ANOVA, pvalues indicated by asterisks with *≤0.05, ***≤0.001, and ****≤0.0001.

## Discussion

Like many other eukaryotic intracellular pathogens, microsporidia transition through distinct stages as they progress through the life cycle. These stages are typically defined through analysis of mixed-stage infections in cell culture and histological sections. In this paper, we used a pulse-chase technique to follow the progression of synchronized infections initiated by a single microsporidia cell in a single host cell of an intact animal from invasion until the final stage of the life cycle. With these data we provide a comprehensive view of the growth dynamics for microsporidia in a natural niche, and identify a novel mechanism for the intercellular spread of eukaryotic intracellular pathogens.

The most surprising observation that we made with our experimental approach is that microsporidia spread between host cells by fusing them together into syncytia. Cell-cell fusion is a widespread phenomenon in animal development. *C. elegans* exemplifies this, with more than 30% of all somatic nuclei sharing cytoplasmic space [27]. To our knowledge, this is the first description of a eukaryotic pathogen causing syncytia formation in animal host tissues. Viruses and bacteria have been shown previously to spread by causing host cell fusions in several animals, including a virus that infects and causes syncytia formation in the intestine of *C. elegans* [28]. Structural and mechanistic distinctions can be made between enveloped viruses that cause host cell fusion through expression of viral entry receptors on the host cell surface that end up binding to and fusing with uninfected cells (e.g. influenza, HIV, and ebola viruses), and non-enveloped reoviruses that fuse host cells solely to spread from an infected into an uninfected host cell [29]. The structural and molecular mechanisms underlying bacterial pathogen-induced host cell fusion have not yet been identified. We do not yet know if there are any structural similarities between the canonical host cell fusion processes and the microsporidia-induced host cell fusions that we describe here. As such, we use the term ‘fusion’ as a simple way to describe the joining together of cells that are normally separated by membranes. It remains a possibility that *Nematocida* microsporidia cells spread to span several neighboring *C. elegans* cells by breaking down their lateral membranes as opposed to fusing the two lipid bilayers. Further studies on the molecular and structural changes involved in the *Nematocida*-*C. elegans* interactions we described here will contribute to our understanding of how two separate cells can be joined together.

We observed species-specific patterns in the host cell fusions induced by the microsporidian pathogens of *C. elegans*. Specifically, *N. parisii* and *N*. sp. 1 caused intestinal host cell fusions but did not grow in other tissues, while *N. displodere* grew in several tissues causing host cell fusions in the muscle and epidermis but not the intestine. The failure of *N. displodere* to spread across intestinal cells was not due to an inability to grow within the tissue, as single intestinal cells were filled with pathogen that differentiated and completed its life cycle. These patterns suggest that there is variation in the factors used by *Nematocida* to cause host cell fusion in *C. elegans*. The general consistency of *Nematocida-*induced syncytia formation in distinct tissues raises the question of why a pathogen would evolve to grow by causing host cell fusion. A common hypothesis for why intracellular pathogens remain in the host cytoplasm during cell-cell spread is to limit the exposure to potentially harmful factors in the extracellular space, such as immune cells. While *C. elegans* does not have any known professional immune cells, other extracellular factors such as antimicrobials including C-type lectins, lysozymes and lipases might be detrimental to microsporidia in this host. Another possibility involves the rate of access to host resources. In the case of microsporidia, we have previously shown that *N. parisii* can escape host intestinal cells by differentiating into spores and exiting through the exocytic pathway [30]. Spores are specialized structures that can withstand harsh environmental conditions but must invade host cells and progress through a lag phase before growth. These limitations imply that microsporidia growth might occur more rapidly by accessing additional host space and resources through spreading before differentiating into spores. The shared features of intercellular spread among species of *Nematocida* also raises the question of how conserved this growth strategy is across microsporidia. There have been previous reports on infections of both vertebrates and invertebrates where microsporidia caused the formation of xenomas, which are described as hypertrophic host cells that increase in size and number of nuclei [31–33]. Given the potential difficulties in distinguishing hypertrophy from cell fusion events in histological sections, we speculate that some of the host cell changes in these microsporidia infections may reflect the kind of host cell syncytia formation that we observed for *Nematocida* during intercellular spread. Identifying the factors involved in *Nematocida*-induced host cell fusion will allow us to search for factors that may be important for other microsporidia-host cell interactions.

The experimental approaches taken in this paper allowed us to identify several features of *Nematocida* growth. Despite the lower temperatures used in this study, we measured a similar *N. parisii* doubling time compared to our previous estimates at a higher temperature (2.4 h at 20°C vs 2.9 h at 25°C, respectively) [20]. Thus, microsporidia grow faster than previously estimated, which may be accounted for in part by differences between the two studies in the synchronicity of infections. Additionally, we found that the doubling rate was uneven over time. The changes in the rate of growth occurred over distinct phases with similarities to other microbes including a lag phase without replication, exponential growth and replication phase, and a slowing to stationary phase at the time of spore differentiation. There are several definitions for the lag phase of microbial growth, but it can be generally described as the time it takes for a microbe to begin growing in biomass and replicating after inoculation in growth media [34]. A recent study identified transcriptional signatures of two distinct sub-phases in the *E. coli* lag phase: one in which there is metabolic activity without an increase in biomass, and a second in which there is an increase in biomass but not yet cell number [35]. While there have been advances in our understanding of the lag phase for bacteria, virtually nothing is known about the lag phase for eukaryotic pathogens. As the timing of the lag phase of growth can profoundly alter the outcome of interactions with a host [36], it will be important to determine how comparable the lag phases are among pathogenic microbes. Studying the lag phase of obligate intracellular pathogens like microsporidia will require strategies for overcoming the challenge of extracting signal from the extremely low ratios in pathogen to host material during this stage of infection.

Genetic variation can be a strong determinant of host-pathogen interactions [37]. We previously showed that there is variation among *C. elegans* strains in their resistance to microsporidia infection, which is a complex genetic trait [21]. Here we show that there is genetic variation among microsporidia in their ability to grow and affect the fitness of *C. elegans*. Through our observations of the differences between microsporidia species in their rates of growth, we were pointed towards factors that influence the kinetics of the microsporidia life cycle. We found that increasing the density of infection by decreasing the host body size or increasing the number of infectious events sped up the development of spores. Neither of these modifications changed the growth rate of *N. parisii*, indicating that the transition from replication to spore differentiation is regulated by environmental factors in the host. Our experiments lead us to speculate that the cue for transitioning from replication to spore formation in microsporidia may be related to the sensing of host resource availability and/or the sensing of self, both of which are related to the density of infection. There is precedent for eukaryotic organisms making developmental decisions based on density, which is exemplified by several fungal species that undergo developmental switches in response to high density and nutrient availability [38]. As has been shown for many other developmental switches in pathogens [39], the specific factors that regulate transitions in the microsporidia life cycle likely involve responses made by both host and pathogen to changes in their shared metabolic pool. Our finding that microsporidia can grow across a substantial portion of the cells comprising an entire animal organ before differentiating opens up a new set of questions regarding the dynamic inputs that intracellular pathogens interpret from the host environment to optimize growth and transmission.

## Materials and methods

### *C. elegans* and *Nematocida* strains

*C. elegans* strains were maintained on nematode growth media (NGM) seeded with *E. coli* OP50-1 (which is a streptomycin-resistant OP50 strain) as previously described [40]. For simplicity, this strain is referred to as OP50 throughout. To obtain starved and synchronized L1 larvae, gravid adults were bleached to isolate eggs, which then were allowed to hatch overnight at 20°C [41]. The *C. elegans* wild-type N2, small mutant CB1482 *sma-6(e1482)*, long mutant CB185 *lon-1(e185)*, and intestinal GFP transgenic SJ4144 *zcIs18 [ges-1p::GFP(cyt)]* strains were obtained from the Caenorhabditis Genetics Center. ERT351 was derived from SJ4144 by backcrossing to N2 eight times. Transgenic animals with intestinal expression of GFP-labeled LET-413 were generated by injection of pET213 *[vha-6p::GFP::let-413]* and a *myo-2::mCherry* co-injection marker to generate a multi-copy array strain. This strain was treated with UV psoralen at 700μJ to generate the integrated ERT147 *jyIs21[vha-6p::GFP::let-413, myo-2::mCherry]* strain. A cytoplasmic intestinal photoconvertible fluorescent protein construct pET207 was generated using three-part Gateway recombination by fusing the intestinal-specific *vha-6* promoter to Dendra [42] with the *unc-54* 3’UTR. This construct was injected into N2 animals and transgenic progeny were recovered to generate a multi-copy array strain ERT113 *jyEx46[vha-6p::dendra::unc-54 3’UTR]*. Infection experiments were performed with *Nematocida parisii* strain ERTm1, *Nematocida* sp. 1 strain ERTm2, and *Nematocida displodere* strain JUm2807 [17,18]. Spores were prepared and quantified as previously described [43].

### Single microsporidia cell infections

Synchronized first-larval stage (L1) animals were inoculated with OP50 and a series of spore dilutions on NGM plates at 20°C. These animals were collected two and a half hours after plating, washed three times with PBS containing 0.1% Tween 20 to remove spores, and re-plated with OP50 at 20°C. A fraction of the population was fixed at this time with 4% PFA for 30 minutes, then stained by FISH with the *Nematocida* ribosomal RNA-specific MicroB probe [18] conjugated to a red Cal Fluor 610 dye (Biosearch Technologies). The number of infectious events per animal was quantified in 50 animals per dosage after mounting samples on agarose pads with VECTASHIELD mounting medium containing DAPI (Vector Labs) and imaging using a Zeiss AxioImager M1 upright fluorescent microscope with a 40X oil immersion objective. A limiting dilution of *Nematocida* spores was tested to identify a concentration in which more than 83% of infected animals contained only a single microsporidia cell. The spore dosage that had this characteristic typically yielded a Poisson distribution of infection, in which an average of 72% of the population was uninfected and 28% infected. Infection distributions were measured for all experiments to ensure that the vast majority of infected animals contained a single microsporidia cell (see Table S1).

### Measuring microsporidia growth by microscopy

The ERT351 strain was infected at the L1 stage at a dosage following the parameters described above for single microsporidia cell infections. Animals were fixed at various times post-inoculation in 4% PFA for FISH staining and microscopy-based analysis of growth. Samples were mounted on 5% agarose pads and imaged using a 40X oil immersion objective or a 10X objective on a Zeiss LSM700 confocal microscope run by ZEN2010 software. Z-stacks were acquired of the entire intestinal space with a z-spacing of 1μm (at 400X) or 6μm (100X), collecting GFP signal (expressed in cytoplasm of intestine) and RFP signal (microsporidia signal stained by FISH). Analysis of these images was performed with Fiji software [44]. Briefly, the brightest slice was used to threshold GFP and RFP to binary signals. The 3D objects counter function was used to measure voxels after converting signals, which were used to quantify the volume of intestine (GFP) and pathogen (RFP) in 10 animals per time point.

### Analysis of microsporidia growth across host intestinal cells

ERT147 animals were infected with a single microsporidia cell. These animals were imaged at various times post-inoculation either after being fixed and stained as described above or as live animals using a 40X oil immersion objective on a Zeiss LSM700 confocal microscope run by ZEN2010 software. For live imaging, infected ERT147 animals were mounted on 5% agarose pads with microsphere beads and imaged with a 63X oil-immersion objective for 120 cycles at 1.58 μs pixel dwell time for five minutes. GFP was excited with a 488nm laser at 2.1% power, and light was collected with the pinhole set to 44μm. Quantifying growth of microsporidia across intestinal cell boundaries was performed after fixing ERT147 animals and staining by FISH as described above. To measure the diffusion of cytoplasm in intestinal cells, ERT113 animals were infected at the L1 stage and imaged live 48 hours post-inoculation with a 40X oil-immersion objective. A 488 nm laser at 0.5% power was used to excite green Dendra proteins and a 555 nm laser at 14% power was used to excite red Dendra proteins. Light was collected with a pinhole size of 42 μm and a 3.15 μs pixel dwell time. Dendra proteins were converted in a 4 μm diameter space with a 405 nm laser at 35% power with a pixel dwell time of 50 μs. Infected cells could be distinguished from uninfected cells based on exclusion of fluorescent signal. To quantify the diffusion of converted signal, the mean red signal was measured in a 15 μm^2^ area within the targeted cell or outside of the targeted cell (at a distance of greater than 30 μm from targeted cell) before and two minutes after conversion.

### Comparing *Nematocida* growth across different host tissues

Transgenic animals expressing GFP in the cytoplasm and nuclei of intestinal or muscle cells were infected with single *N. parisii, N*. sp. 1, or *N. displodere* cells at 15°C and fixed at the time of spore formation. For the intestinal-GFP strain ERT413 *jySi21[spp-5p::GFP; cb-unc-119(*)] II* [17], 1000 synchronized L1 larvae were grown at 20°C for 48 hours to the young adult stage before being inoculated with 3.7 x 10^5^ *N. parisii* spores, 3.2 x 10^5^ *N*. sp. 1 spores, or 1.25 x 10^4^ *N. displodere* spores for 30 minutes. Spores were washed off and infection progressed at 15°C. For the muscle-GFP strain HC46 *ccIs4251[myo-3::GFP-NLS, myo-3::GFPMITO] I; mIs11[myo-2::GFP] IV* [45], 1000 synchronized L1 larvae were infected in duplicate at 15°C with 5.0-20.0 x 10^4^ spores of *N. displodere* on a 6 cm plate. A fraction of animals was fixed in 4% PFA diluted in PBS * 0.1% Tween-20 at 24 hpi for FISH to verify infection distributions. At 76 hpi (*N. parisii* and *N*. sp. 1) or 120 hpi (*N. displodere*), animals were fixed for FISH with or without DY96 (10μg/ml) staining to stain infection and spores. *N. displodere* FISH was performed as previously described [17]. All analysis and imaging was conducted on a Zeiss LSM700 confocal microscope with a 40x oil-immersion objective. The numbers of GFP* host cell nuclei abutted by pathogen in a 3D stack of confocal images were counted to quantify the number of host cells that infection had spread into by the time of sporulation.

### Measuring microsporidia growth by qPCR

The N2 strain was infected at the L1 stage at a dosage following the parameters described above for single microsporidia cell infections. Animals were fixed at various times post-inoculation in Extracta (Quanta Biosciences) to isolate DNA for qPCR-based analysis of growth. Microsporidia copy number was quantified with iQ SYBR Green Supermix (Bio-Rad) on a CFX Connect Real-time PCR Detection System (Bio-Rad). We measured the relative abundance of *Nematocida* and *C. elegans* DNA in 30 ng of DNA with the following primer sets: Np_rDNAF1: aaaaggcaccaggttgattc, Np_rDNAR1: agctctctgacgcttccttc, Ce_snb-F1: ccggataagaccatcttgacg, Ce_snb-R1: gacgacttcatcaacctgagc. Microsporidia copy number was measured by normalizing to samples infected by single microsporidia cells at early stages of growth when only a single nucleus is observed. We validated qPCR measurements of microsporidia copy number by taking a microscopy-based method of manually counting the average number of microsporidia nuclei per infection during growth from a single nucleus to 100 nuclei. Copy number measurement discrepancies between qPCR and manual count analyses were within 11%. Primer efficiencies were measured, and fold difference was calculated using the Livak comparative Ct method (2-ΔΔCt).

### Measuring animal size, egg number, and microsporidia spores

Animals were fixed and stained by FISH as described above, then imaged using a Zeiss AxioImager M1 upright fluorescent microscope with a 40X oil immersion objective. 50 animals per condition were manually outlined with Fiji software to calculate size (in μm^2^). The number of eggs per animal was measured at 60 hpi by adding DY96 (4ng/ml with 0.1% SDS) to FISH-stained samples for 30 minutes before imaging. DY96 stains chitin [46], which is a component of the eggshell. The 60 hpi time point was chosen because it is when uninfected animals first consistently contain eggs. Eggs were counted in 50 animals per condition. Microsporidia spores also contain chitin, and were stained with DY96. Spore clusters per infected animal in 50 total animals per condition were counted. Each cluster of spores contained approximately 50 spores. Spore clusters (which we define as regions within meronts that contain approximately 30 spores with bright chitin staining) were counted at 52 hpi, which is the time at which single microsporidia cell infections begin to differentiate in to spores in N2 animals as described above.

## Author Contributions

KMB, RJL, and ERT designed experiments, KMB and RJL performed experiments and analyzed data, MAB generated the ERT147 transgenic strain, RJL and MAB contributed to the manuscript, KMB and ERT wrote the manuscript.

## Acknowledgements

We thank Michael Botts, Kirthi Reddy, and Aaron Reinke for helpful comments on the manuscript, and the *Caenorhabditis* Genetics Center for *C. elegans* strains.

## References

1. Bernardini ML, Mounier J, d’Hauteville H, Coquis-Rondon M, Sansonetti PJ (1989) Identification of icsA, a plasmid locus of Shigella flexneri that governs bacterial intra- and intercellular spread through interaction with F-actin. Proc Natl Acad Sci U S A 86: 3867–3871.

2. Heinzen RA, Hayes SF, Peacock MG, Hackstadt T (1993) Directional actin polymerization associated with spotted fever group Rickettsia infection of Vero cells. Infect Immun 61: 1926–1935.

3. Tilney LG, Portnoy DA (1989) Actin filaments and the growth, movement, and spread of the intracellular bacterial parasite, Listeria monocytogenes. J Cell Biol 109: 1597–1608.

4. Cudmore S, Cossart P, Griffiths G, Way M (1995) Actin-based motility of vaccinia virus. Nature 378: 636–638.

5. Ciechonska M, Duncan R (2014) Reovirus FAST proteins: virus-encoded cellular fusogens. Trends Microbiol 22: 715–724.

6. French CT, Toesca IJ, Wu TH, Teslaa T, Beaty SM, et al. (2011) Dissection of the Burkholderia intracellular life cycle using a photothermal nanoblade. Proc Natl Acad Sci U S A 108: 12095–12100.

7. Steele S, Radlinski L, Taft-Benz S, Brunton J, Kawula TH (2016) Trogocytosis-associated cell to cell spread of intracellular bacterial pathogens. Elife 5.

8. Nikitas G, Deschamps C, Disson O, Niault T, Cossart P, et al. (2011) Transcytosis of Listeria monocytogenes across the intestinal barrier upon specific targeting of goblet cell accessible E-cadherin. J Exp Med 208: 2263–2277.

9. Swann J, Jamshidi N, Lewis NE, Winzeler EA (2015) Systems analysis of host-parasite interactions. Wiley Interdiscip Rev Syst Biol Med 7: 381–400.

10. Sturm A, Amino R, van de Sand C, Regen T, Retzlaff S, et al. (2006) Manipulation of host hepatocytes by the malaria parasite for delivery into liver sinusoids. Science 313: 1287–1290.

11. Stentiford GD, Becnel JJ, Weiss LM, Keeling PJ, Didier ES, et al. (2016) Microsporidia - Emergent Pathogens in the Global Food Chain. Trends Parasitol 10.1016/j.pt.2015.12.004.

12. Stentiford GD, Feist SW, Stone DM, Bateman KS, Dunn AM (2013) Microsporidia: diverse, dynamic, and emergent pathogens in aquatic systems. Trends Parasitol 29: 567–578.

13. Cali A, Takvorian PM (2014) Developmental Morphology and Life Cycles of the Microsporidia. Microsporidia: John Wiley & Sons, Inc. pp. 71–133.

14. Xu Y, Weiss LM (2005) The microsporidian polar tube: a highly specialised invasion organelle. Int J Parasitol 35: 941–953.

15. Balla KM, Troemel ER (2013) Caenorhabditis elegans as a model for intracellular pathogen infection. Cell Microbiol 10.1111/cmi.12152.

16. Felix MA, Duveau F (2012) Population dynamics and habitat sharing of natural populations of Caenorhabditis elegans and C. briggsae. BMC biology 10: 59.

17. Luallen RJ, Reinke AW, Tong L, Botts MR, Felix M-A, et al. (2016) Discovery of a Natural Microsporidian Pathogen with a Broad Tissue Tropism in Caenorhabditis elegans. bioRxiv 10.1101/047720.

18. Troemel ER, Felix MA, Whiteman NK, Barriere A, Ausubel FM (2008) Microsporidia are natural intracellular parasites of the nematode *Caenorhabditis elegans*. PLoS Biol 6: 2736–2752.

19. Bakowski MA, Desjardins CA, Smelkinson MG, Dunbar TL, Lopez-Moyado IF, et al. (2014) Ubiquitin-mediated response to microsporidia and virus infection in C. elegans. PLoS Pathog 10: e1004200.

20. Cuomo CA, Desjardins CA, Bakowski MA, Goldberg J, Ma AT, et al. (2012) Microsporidian genome analysis reveals evolutionary strategies for obligate intracellular growth. Genome research 22: 2478–2488.

21. Balla KM, Andersen EC, Kruglyak L, Troemel ER (2015) A wild C. elegans strain has enhanced epithelial immunity to a natural microsporidian parasite. PLoS Pathog 11: e1004583.

22. Legouis R, Gansmuller A, Sookhareea S, Bosher JM, Baillie DL, et al. (2000) LET-413 is a basolateral protein required for the assembly of adherens junctions in Caenorhabditis elegans. Nat Cell Biol 2: 415–422.

23. Altun Z.F. HDH (2009) Muscle system, somatic muscle. WormAtlas.

24. Altun Z.F. HDH (2009) Epithelial system, hypodermis. WormAtlas.

25. Krishna S, Maduzia LL, Padgett RW (1999) Specificity of TGFbeta signaling is conferred by distinct type I receptors and their associated SMAD proteins in Caenorhabditis elegans. Development 126: 251–260.

26. Maduzia LL, Gumienny TL, Zimmerman CM, Wang H, Shetgiri P, et al. (2002) lon-1 regulates Caenorhabditis elegans body size downstream of the dbl-1 TGF beta signaling pathway. Dev Biol 246: 418–428.

27. Podbilewicz B (2006) Cell fusion. In: Community TCeR, editor. WormBook: WormBook.

28. Felix MA, Ashe A, Piffaretti J, Wu G, Nuez I, et al. (2011) Natural and experimental infection of Caenorhabditis nematodes by novel viruses related to nodaviruses. PLoS Biol 9: e1000586.

29. Sapir A, Avinoam O, Podbilewicz B, Chernomordik LV (2008) Viral and developmental cell fusion mechanisms: conservation and divergence. Dev Cell 14: 11–21.

30. Szumowski SC, Botts MR, Popovich JJ, Smelkinson MG, Troemel ER (2014) The small GTPase RAB-11 directs polarized exocytosis of the intracellular pathogen N. parisii for fecal-oral transmission from C. elegans. Proc Natl Acad Sci U S A 10.1073/pnas.1400696111.

31. Leitch GJ, Shaw AP, Colden-Stanfield M, Scanlon M, Visvesvara GS (2005) Multinucleate host cells induced by Vittaforma corneae (Microsporidia). Folia Parasitol (Praha) 52: 103–110.

32. Lom J, Dykova I (2005) Microsporidian xenomas in fish seen in wider perspective. Folia Parasitol (Praha) 52: 69–81.

33. Maurand J (1973) Recherches biologiques sur les microsporidies des larves de simulies. CNRS A: Academie de Montpellier.

34. Monod J (1949) The Growth of Bacterial Cultures. Annual Review of Microbiology 3: 371–394.

35. Madar D, Dekel E, Bren A, Zimmer A, Porat Z, et al. (2013) Promoter activity dynamics in the lag phase of Escherichia coli. BMC Syst Biol 7: 136.

36. Stepanyan K, Wenseleers T, Duenez-Guzman EA, Muratori F, Van den Bergh B, et al. (2015) Fitness trade-offs explain low levels of persister cells in the opportunistic pathogen Pseudomonas aeruginosa. Mol Ecol 24: 1572–1583.

37. Persson J, Vance RE (2007) Genetics-squared: combining host and pathogen genetics in the analysis of innate immunity and bacterial virulence. Immunogenetics 59: 761–778.

38. Sprague GF, Jr., Winans SC (2006) Eukaryotes learn how to count: quorum sensing by yeast. Genes Dev 20: 1045–1049.

39. Olive AJ, Sassetti CM (2016) Metabolic crosstalk between host and pathogen: sensing, adapting and competing. Nat Rev Microbiol 14: 221–234.

40. Brenner S (1974) The genetics of *Caenorhabditis elegans*. Genetics 77: 71–94.

41. Stiernagle T (2006) Maintenance of C. elegans. In: Community TCeR, editor. WormBook: WormBook.

42. Gurskaya NG, Verkhusha VV, Shcheglov AS, Staroverov DB, Chepurnykh TV, et al. (2006) Engineering of a monomeric green-to-red photoactivatable fluorescent protein induced by blue light. Nat Biotechnol 24: 461–465.

43. Estes KA, Szumowski SC, Troemel ER (2011) Non-lytic, actin-based exit of intracellular parasites from *C. elegans* intestinal cells. PLoS Pathog 7: e1002227.

44. Schindelin J, Arganda-Carreras I, Frise E, Kaynig V, Longair M, et al. (2012) Fiji: an open-source platform for biological-image analysis. Nat Methods 9: 676–682.

45. Winston WM, Molodowitch C, Hunter CP (2002) Systemic RNAi in C. elegans requires the putative transmembrane protein SID-1. Science 295: 2456–2459.

46. Hoch HC, Galvani CD, Szarowski DH, Turner JN (2005) Two new fluorescent dyes applicable for visualization of fungal cell walls. Mycologia 97: 580–588.

